# MPBind: Multitask Protein Binding Site Prediction by Protein Language Models and Equivariant Graph Neural Networks

**DOI:** 10.1101/2025.04.12.648527

**Authors:** Yanli Wang, Frimpong Boadu, Jianlin Cheng

## Abstract

Proteins interact with a variety of molecules, including other proteins, DNAs, RNAs, ligands, ions, and lipids. These interactions play a crucial role in cellular communication, metabolic regulation, immune response, and structural integrity, making proteins fundamental to nearly all biological functions. Accurately predicting protein binding (interaction) sites is essential for understanding protein interactions and functions. Here, we introduce MPBind, a multitask protein binding site prediction method, which integrates protein language models (PLMs) that can extract structural and functional information from sequences and equivariant graph neural networks (EGNNs) that can effectively capture geometric features of 3D protein structures. Through multitask learning, it can predict binding sites on proteins that interact with five key categories of binding partners: proteins, DNA/RNA, ligands, lipids, and ions, which is more comprehensive than most existing task-specific methods that can predict only one or a few kinds of binding sites. Moreover, MPBind outperforms both general bind site prediction methods and task-specific binding site prediction methods.

## 1. Introduction

Proteins are highly versatile biomolecules that interact with a wide range of molecules, including other proteins, DNA, RNA, ligands, ions, and lipids, to perform essential cellular functions [1]. Rather than working alone, they function as part of larger molecular machines or complexes, where their roles depend on the specific interactions they form [2]. These interactions are essential for key cellular processes like metabolism, DNA replication, gene expression, signaling, and structural support [3]. A protein’s ability to bind with other molecules comes from its unique three-dimensional shape and specialized features, such as domains and motifs, which help it make precise interactions [4]. Understanding how proteins interact not only offers valuable insights into cellular mechanisms but also helps guide the development of therapeutic strategies by leveraging these core biological processes. With the explosive growth of protein sequences and structures, accurately predicting protein binding sites (i.e., the specific residues of a protein that interact with its partners) is essential for decoding protein interactions and their biological roles, which is important for drug design and disease treatment.

The rapid increase in experimental and predicted protein structures has created both a major challenge and an opportunity for using this structural information to represent and predict protein surface properties such as binding sites [5]. In recent years, deep learning has greatly changed the game when it comes to predicting protein binding sites [6]. Deep learning models can now process huge amounts of protein structure data, automatically identifying complex patterns and predicting binding sites with better accuracy than before [6, 7]. Based on the input information used by deep learning methods, they can be largely grouped into two categories: sequence-based [7] and structure-based [6] methods. Structure-based approaches typically focus on using the three-dimensional (3D) structure of proteins to pinpoint where binding sites are located, taking advantage of the arrangement of atoms and the overall protein shape to identify potential binding sites [6]. In this category, deep learning methods typically represent 3D structures using grid-like or voxel-based models [8-10], as well as graph-based representations [11-14], to detect patterns and identify regions likely to interact with other molecules, such as ligands, other proteins, or DNA. Unlike structure-based methods, sequence-based deep learning approaches predict protein binding sites using only the amino acid sequence, without leveraging the protein’s 3D structure [7]. These methods rely on deep learning models to recognize patterns in protein sequences and pinpoint potential binding sites [15-18], making them useful for proteins with no known structural data. For instance, CLAPE[15]takes protein sequences as input to predict protein-DNA binding sites, leveraging a pre-trained protein language model (PLM) and contrastive learning without considering 3D structural information.

As the structures of most proteins can be readily obtained by high-accuracy protein structure prediction methods such as AlphaFold [19, 20], it is important to fully harness the valuable insights from both protein sequences and structures to improve protein binding site prediction. ScanNet [14] takes protein 3D structures and their corresponding sequences as input to predict protein-protein interaction interfaces, primarily emphasizing the geometric information of the structures in alignment with the sequences. However, it does not fully exploit sequence information from pre-trained protein language models (PLMs). PeSTo [11] primarily uses the atomic coordinates of protein structures as input to predict multiple types of binding sites through a geometric transformer. However, like ScanNet, it does not fully utilize sequence information in PLMs. GraphBind [16] leverages both protein structural information and corresponding sequence information through a hierarchical graph neural network. However, it relies on traditional sequence alignment methods, such as HHblits profile [21] and PSI-BLAST profile [22], to explore sequence information. GPSite [17] takes protein sequences as input to predict multiple types of binding sites at the residue level via a graph transformer and leverages the structural information in the structures predicted by ESMfold [23], but it does not utilize more accurate structures predicted by AlphaFold. Therefore, there is a need to fully harness the valuable insights from both protein sequences and structures to further improve protein binding site prediction.

Nowadays, protein language models (PLMs) [24-26], deep learning models designed to understand and analyze protein sequences in a manner similar to how natural language models process human language, have proven to be incredibly effective at analyzing and predicting protein structures, functions, and interactions. By recognizing patterns in vast amounts of protein sequence data, they can be used to identify binding sites, classify protein functions, and even generate new protein sequences. Meanwhile, Equivariant Graph Neural Networks (EGNNs) are an emerging geometric deep learning approach suitable for protein structure analysis, as they can maintain consistent predictions when the input data undergoes transformations such as rotations or translations [27, 28]. Unlike traditional graph neural networks (GNNs), which operate on 2D graph structures (nodes and edges) without considering 3D locations, EGNNs offer a robust framework for analyzing and predicting protein structures and interactions by leveraging symmetry and 3D spatial information [12, 28]. Therefore, it is interesting to leverage the strengths of both PLMs and EGNNs to improve protein binding site prediction, especially for multitask predictions across different types of binding sites.

In this work, we introduce MPBind, a multitask deep learning method to predict five kinds of binding sites through one model. MPBind combines two protein language models [25, 26], pre-trained on different datasets to provide complementary insights, with an equivariant graph neural network [28] to capture and process both sequence and structural information. Additional information, such as secondary structures, atomic and geometric residue-level details, and geometric edge information, is also incorporated into the model. Using a multitask framework, MPBind can predict and analyze a wide range of protein binding sites, including interactions with other proteins, DNA, RNA, ligands, ions, and lipids. After training on experimental structures from the PDB database [29, 30], it can predict binding sites from either experimental structures or AlphaFold-predicted structures (e.g., ones in AlphaFold Protein Structure Database [20, 31]).

Compared to other deep learning models designed for both task-specific and general binding site predictions, MPBind outperforms general (multitask) prediction methods and achieves competitive performance with the best task-specific method. Further analysis of predicted binding sites in human proteomics data, combined with UniProt-annotated features [32], indicates that the predictions by MPBind can reveal important links between binding sites, molecular functions, and biological processes.

## 2. Methods

### 2.1 Datasets for training, validating, and testing MPBind

Two datasets are used in this work. The first dataset (Dataset1) comes from PeSTo [11], which includes the protein structures released before January 1, 2022. Following the exact data processing procedure of PeSto, we downloaded all the protein structures from the Protein Data Bank (PDB) [29, 30], extracted all the chains (subunits) of the structures, and clustered them based on a maximum sequence identity of 30%. The clustered chains were partitioned into training, validation, and test sets according to the exact same split used by PeSTo [11] such that the proteins between different sets have less than 30% sequence identity. The three subsets are referred to as ***training_data***, ***validation_data***, and ***Test_data***, which has 376,216 chains (∼70% of the data), 101,700 chains (∼15%), and 97,424 chains (∼15%), respectively. Because ***Test_data*** is too large for any method to run efficiently, we selected all the chains from it using a threshold of a maximum of 8,192 atoms (1,024 × 8) and a minimum of 48 residues, resulting in 40,651 chains. They form a test set called ***Test1_data*** to compare MPBind and PeSTo.

We downloaded the protein structures released after January 1, 2022 and before June 21, 2024, to create the second dataset (Dataset2). The chains in Dataset2 that have >30% sequence identity with the proteins in Dataset1 were removed. The remaining chains were clustered according to the 30% sequence identity threshold. One chain was selected from each cluster to form a set of test proteins, referred to as ***Test2_data***, which contains 1,452 chains. Since the proteins in ***Test2_data*** were released after the latest release date of the training/validation proteins of all the methods evaluated in this work and have <= 30% sequence identity with them, it is new for all the methods and can therefore serve as an objective and rigorous benchmark for them.

To further validate and analyze MPBind, we also downloaded the 23,391 human protein structures in the human proteome from the AlphaFold-European Bioinformatics Institute (AF-EBI) database [20, 31]. This set is used to investigate how well MPBind works on predicted protein structures and how its predictions can be used to study biological functions.

### 2.2 Protein Structure Preprocessing and Label Extraction

Following the same procedure of PeSTo [11], in a protein structure, deuterium atoms, hydrogen, water, and heavy water are all removed. Non-native molecules used to assist in solving the structures are also excluded. If an atom has alternate locations, only the first one is retained.

In a protein structure, a total of 79 kinds of molecules, including 20 standard amino acids, 8 most common nucleic acids, 16 most common ions, 31 most common ligands, and 4 most common lipids, are selected. Two molecules are considered in contact/interaction if the minimum distance between their heavy atoms is within 5 Å. The residues in a chain that are in contact with other types of molecules (nucleotides, ions, ligands, lipids) or residues in other chains are defined as the interface (binding sites). Based on the threshold, a 79×79 interface-type matrix is constructed for all interactions among these molecules, covering protein, DNA/RNA, ion, ligand, and lipid interactions. However, since we focus only on interactions between 20 standard amino acids (residues) of a protein and other molecules, the interface type matrix is reduced to 20×79. In this matrix, entries marked with 1 indicate binding sites, while entries marked with 0 indicate non-binding sites. For a protein chain with *L* residues interacting with another molecule (a protein chain, DNA/RNA, ion, ligand, or lipid) of length M, there are *L* × *M* pairwise combinations that may or may not be in contact. Each combination is projected onto a corresponding interface type matrix of shape 20 × 79, leading to a chain-specific interaction interface matrix of shape *L* × *M* × *20* × *79*, which serves as label for training, validation, and evaluation of MPBind.

### 2.3 Feature Extraction

MPBind uses a graph to represent the input structure of a protein chain and its corresponding sequence information, where a node denotes a residue and an edge connecting two residues (nodes) if the distance between their *C*_*a*_ (alpha carbon) atoms is within 15 Å. Two types of features, node features and edge features, are extracted to represent a protein, as shown in **Fig. 1**.

**Fig. 1:**
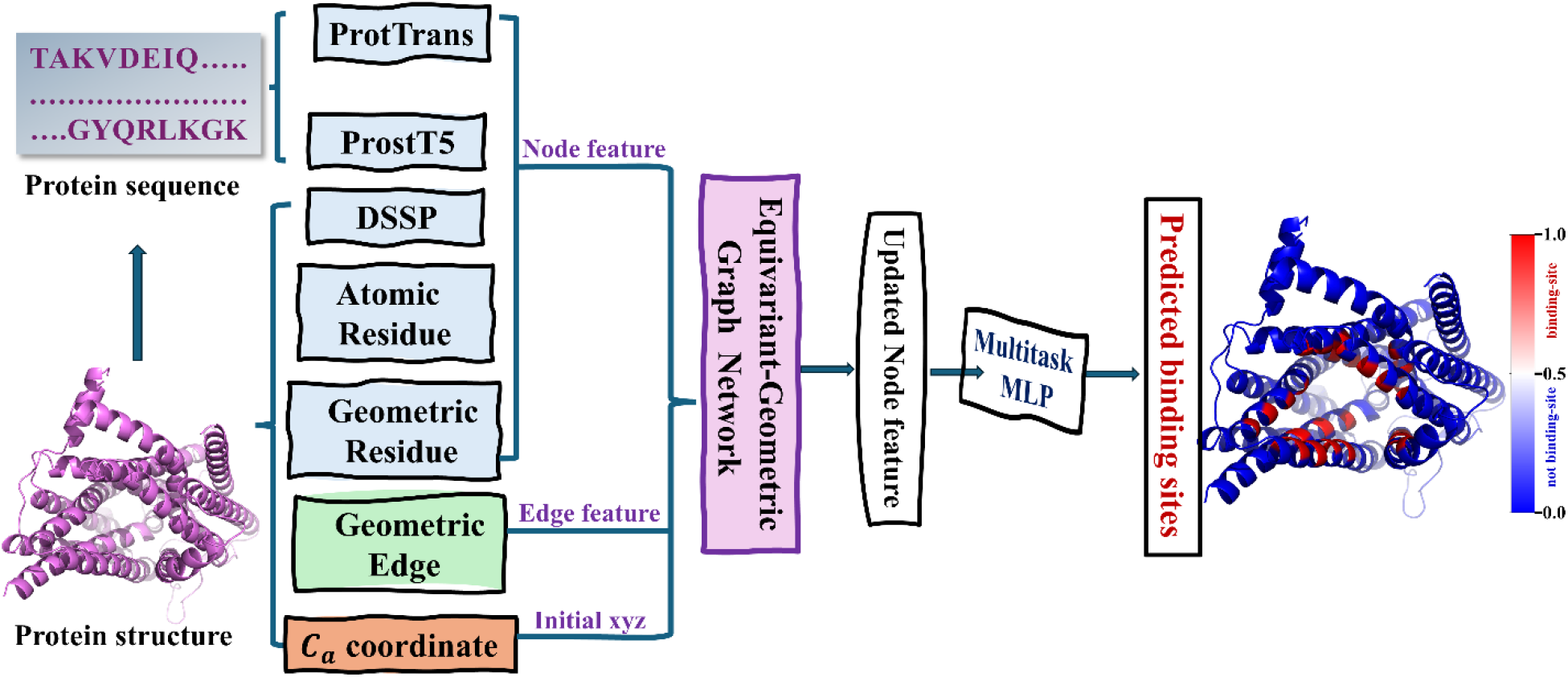
The overall architecture of MPBind. Both protein sequence and structure are used to generate node features from different perspectives. The protein sequence is processed through two PLMs, ProtTrans and ProstT5, to extract residue embeddings. The protein structure provides secondary structure node features extracted by DSSP, atomic residue features derived from intra-residue atomic properties, and geometric residue features obtained from intra-residue atomic coordinates. Additionally, geometric edges are extracted from the protein structure. The five types of node features, along with the geometric edge features and the x,y,z coordinates of Ca atom of each node, are used as inputs to a four-layer equivariant graph neural network block to update node features. The updated node features are then passed through a multitask multi-layer perceptron (MLP) to generate a predicted score ranging from 0 to 1 for each node and each type of binding site. Each score for a node is the predicted probability that the node is a binding site of a specific type.

To fully leverage the node information, we introduce five types of node features extracted from both protein sequences and structures, each of which makes a significant contribution to our model according to an ablation study, which are described as follows.

Both the first and second types of node features are based on the embedding information extracted from protein sequences by Protein Language Models (PLMs) [18, 25, 26], as they have shown better performance in various protein property prediction problems than traditional approaches of using multiple sequence alignment (MSA) as input. Two pre-trained PLMs: ProtTrans [26] and ProstT5 [25], which were trained on different datasets and offer complementary insights, are incorporated into our model to extract residue information. ProtTrans is based on a single modality – protein sequence, while ProstT5 is a multimodal PLM considering both sequence and structure. Each PLM provides residue features of size *L* × 1024 for a protein of length *L*, where 1024 denotes the dimension of the features per residue.

The third type of node feature is secondary structures and solvent accessibility, extracted using DSSP [33, 34] from the 3D structures. This approach generates residue-level node features of size *L* × 9 for a protein of length *L*, where the 9 dimensions of the features per residue include the relative solvent accessibility (the solvent accessible surface of a residue normalized by the maximum solvent accessible surface area of its corresponding amino acid type) and the one-hot encoding of the eight possible secondary structure types.

The fourth type of node feature is the atomic features for each residue. Each residue contains various atoms, and each atom within a specific residue has several properties, such as atomic mass, whether it is a side-chain atom, whether it is part of a ring, the amount of electronic charge, the number of hydrogen atoms bonded to it, and the length of the van der Waals radius. We select these six properties of each atom in a residue as the atomic features. However, since the number of atoms varies across different types of residues, we average the atomic properties within each residue, resulting in atomic node features of size *L* × 6 for one query 3D structure with *L* residues.

The final type of node feature is geometric node features for each residue. Each atom in a residue has a unique position in a protein structure, and the side-chain atoms of a residue can be centralized by averaging their positions, denoted as R. Therefore, the position of each residue can be represented by the x, y, and z coordinates of four main backbone atoms (*N, C, C*_*a*_, *O*) and *R*. Using this position information, we can calculate the bond and torsion angles, intra-residue distances, and relative directions of other inner atoms (i.e., *N, C, O*, and R) to atom *C*_*a*_ respectively, which form the geometric information for each residue (i.e., geometric residue). These resulting geometric node features have a size of *L* × 184 for a protein structure with *L* residues.

Finally, the five types of node features above are concatenated to form the final node features, resulting in a size of *L* × 2247 for an input protein with *L* residues.

To extract edge features, we treat each residue as a group of five (virtual) atoms: *N*, *C*, *C*_*a*_, *O*, and *R*, where R is a virtual atom denoting the centroid of the side chain. We explore several pieces of geometric edge information, including the spatial orientation, calculated by 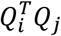, where *Q*_*i*_ denotes the coordinates of *i*-th residue, between adjacent residues connected by an edge, the relative directions of all atoms in a neighboring residue to the *C*_*a*_ atom of the residue in consideration, and the distances between atoms from two residues connected by an edge. To obtain additional geometric information about the edge, the positional embedding information is also calculated based on the index difference between two residues, using *C*_*a*_ as a reference. Finally, the geometric edge features include orientation, direction, distance, and positional embedding information, which together form the edge features with a size of *E* × 450, where *E* denotes the number of edges in an input structure.

### 2.4 The architecture of MPBind

MPBind is an equivariant graph neural network (EGNN) [27, 35] that takes the (x, y, z) coordinates of the Ca atom of each node, along with node and edge features, as input. The reason for using only the position of the Ca atom of each node to represent its position in a protein structure is that the other atoms of the node have been encoded in the node features. Moreover, all atoms of each residue as input would make our model cumbersome and complicated because the number of atoms of different types of residues varies.

As shown in **Fig. 1**, the main component of MPBind is the four-layer equivariant graph neural network (EGNN) block, which is similar to the network used in chromosome structure modeling [36]. Each EGNN layer updates the features of each node considering its own features, the messages from its neighboring nodes, and the relative positions between it and the neighboring nodes. The major advantage of the EGNN layer is that its feature update is equivariant to the rotation and translation of protein structure such that it can focus on extracting essential features relevant to prediction tasks. After the four EGNN layers, the updated node features are passed through a multitask multi-layer perceptron (MLP) head, followed by a sigmoid activation function to predict the probability that each node is a binding site for each of the five types of binding partners (proteins, DNA/RNA, ligands, lipids, and ions), respectively.

### 2.5 Training, validation, and Evaluation

The graph representations of all the protein chains in the ***training_data*** dataset were used to train MPBind. The Binary Cross-Entropy loss function was employed to measure the difference between the predicted scores and binding site labels. During training, the loss function of MPBind was minimized using the Adam optimizer with a learning rate of 0.001 to adjust its weights. The ***validation_data*** dataset was used to monitor MPBind’s performance during the training and select the best model based on the lowest validation loss. After training, the selected best MPBind model was blindly tested on two datasets, ***Test1_data*** and ***Test2_data***, for a fine-grained comparison with other methods. Finally, the selected MPBind model was used to predict the interfaces for the proteins in the human proteome dataset [31], whose structures were predicted by AlphaFold2 [18], for further analysis of interface features.

In line with previous works [11, 16, 17], three common metrics, Receiver Operating Characteristic Area Under the Curve (ROC AUC), Precision-Recall Area Under the Curve (PR AUC), and Accuracy (Acc) are selected to evaluate the interface prediction for MPBind and other methods [11, 14-16, 37]. Receiver Operating Characteristic Area Under the Curve (ROC AUC) is a performance metric used in binary classification to assess a method’s ability to distinguish between positive and negative classes at different thresholds. Precision-Recall Area Under the Curve (PR AUC) measures the alignment of predictions with the known gold standard, making it a suitable evaluation metric for highly imbalanced datasets like the ones in this project, where only a small portion of residues of the proteins in the test datasets are binding sites. It is worth noting, ROC AUC and PR AUC, are most important metrics for evaluating the methods across the different test datasets because they are more comprehensive than the other three metrics and are more suitable for the imbalanced test data.

Accuracy (Acc) measures the proportion of correctly classified instances, meaning the number of instances that were accurately predicted, over the total number of instances in the dataset, indicating the model’s overall classification accuracy, and can be calculated by the following equation:

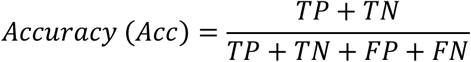

Where TP (True Positive) refers to instances correctly predicted as positive, TN (True Negative) to those correctly predicted as negative, FP to instances incorrectly predicted as positive, and FN to those incorrectly predicted as negative. For highly class-imbalanced datasets with only a small portion of positive labels as in this work, ACC can be very high but still fail to accurately measure how a method distinguishes positive examples from negative ones.

## 3. Results

### 3.1 Protein-protein interface (binding site) prediction

For protein-protein interaction site (binding site) prediction, we first benchmarked MPBind alongside another multitask deep learning binding site prediction method PeSTo [11] on the ***Test1_data*** dataset. Both MPBind and PeSTo are general, non-task-specific prediction methods that can predict different kinds of bind sites. Both of them were trained on the same dataset ***training_data***. Then, we evaluated MPBind, PeSTo, and ScanNet [14] on the ***Test2_data*** dataset. ScanNet is a task-specific geometric deep learning method for predicting protein-protein binding sites, including antibody-antigen binding sites.

The ROC and Precision-Recall curves of MPBind and PeSTo on ***Test1_data*** are presented in **Fig. 2 (A)** and **(B)**. MPBind achieves ROC AUC and PR AUC scores of 0.83 and 0.54, respectively, substantially higher than 0.76 and 0.38 of PeSTo. The results in terms of Accuracy are provided in supplementary **Table S1**, which also shows MPBind outperforms PeSTo. Because MPBind and PeSTo were trained, validated, and tested on the same data, the results demonstrate that MPBind is more accurate and robust than PeSTo.

**Fig. 2:**
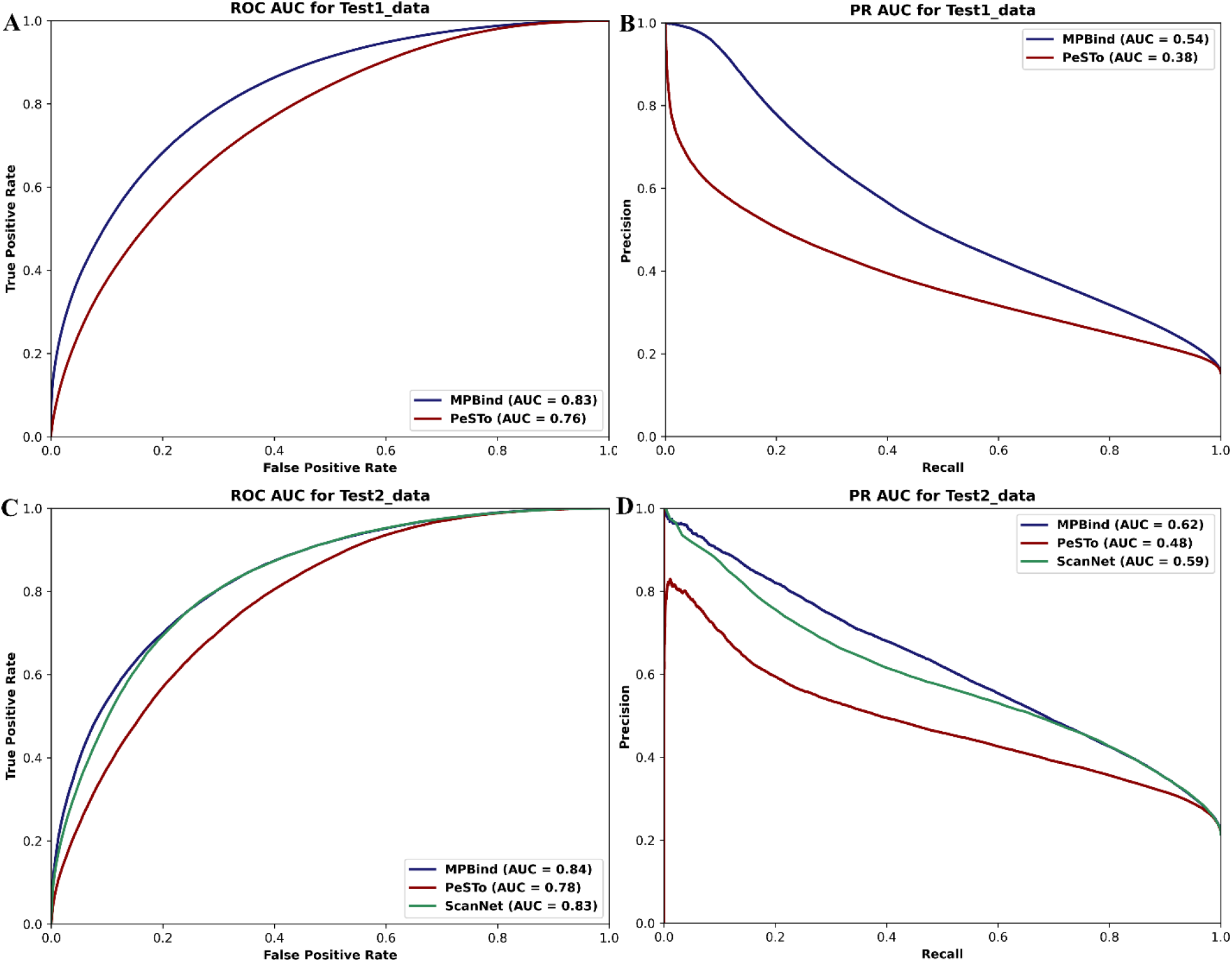
Performance comparison of MPBind and the other two state-of-the-art protein-protein binding site prediction methods across two test datasets. (**A**) ROC curves of MPBind and PeSTo on ***Test1_data*** dataset. (**B**) Precision-Recall curves of MPBind and PeSTo on ***Test1_data*** dataset. (**C**) ROC curves of MPBind, PeSTo, and ScanNet on ***Test2_data*** dataset. (**D**) Precision-Recall curves of MPBind, PeSTo, and ScanNet on ***Test2_data*** dataset. The AUC scores under ROC and PRC curves of the methods are listed in the legend of each sub-figure.

Among the 1,452 chains in ***Test2_data***, 1,208 are involved in protein-protein interactions. Therefore, we compared the task-specific model, ScanNet, and two non-task-specific models, MPBind and PeSTo, for protein binding site prediction on these 1,208 chains. The results for the two critical metrics, ROC AUC and PR AUC, are presented in **Fig. 2 (C)** and **(D)**. Compared to the non-task-specific model PeSTo, MPBind achieves ROC AUC and PR AUC scores of 0.84 and 0.62, respectively, substantially outperforming PeSTo with scores of 0.78 and 0.48. Even when compared to the task-specific model ScanNet dedicated to protein-protein binding site prediction, MPBind still achieves better results, surpassing ScanNet by 0.01 and 0.03 in terms of ROC AUC and PR AUC, respectively. The results in terms of Accuracy are provided in supplementary **Table S1**, where MPBind achieves the highest Accuracy. Overall, these results demonstrate that MPBind achieves state-of-the-art performance in protein-protein binding site prediction.

### 3.2 Robust generalization for DNA/RNA, ion, ligand, and lipid binding site prediction

To evaluate MPBind’s performance in predicting the binding sites for nucleotides (DNA or RNA), ions, ligands, and lipids, we compared it with the same non-task-specific method, PeSTo [11], along with the three top task-specific prediction methods—CLAPE [15], LMetalSite [37], and GraphBind [16]—on the ***Test2_data*** dataset. The dataset includes 75 DNA-binding chains, 48 RNA-binding chains, 406 ion-binding chains, 448 ligand-binding chains, and 39 lipid-binding chains. Notably, some chains may contain multiple binding interfaces. Following the convention in the field [11], the DNA and RNA binding sites are treated as the same type of binding sites (nucleotide binding sites). The results for the two primary metrics, ROC AUC and PR AUC, across four different types of binding site predictions are presented in **Fig. 3**.

**Fig. 3:**
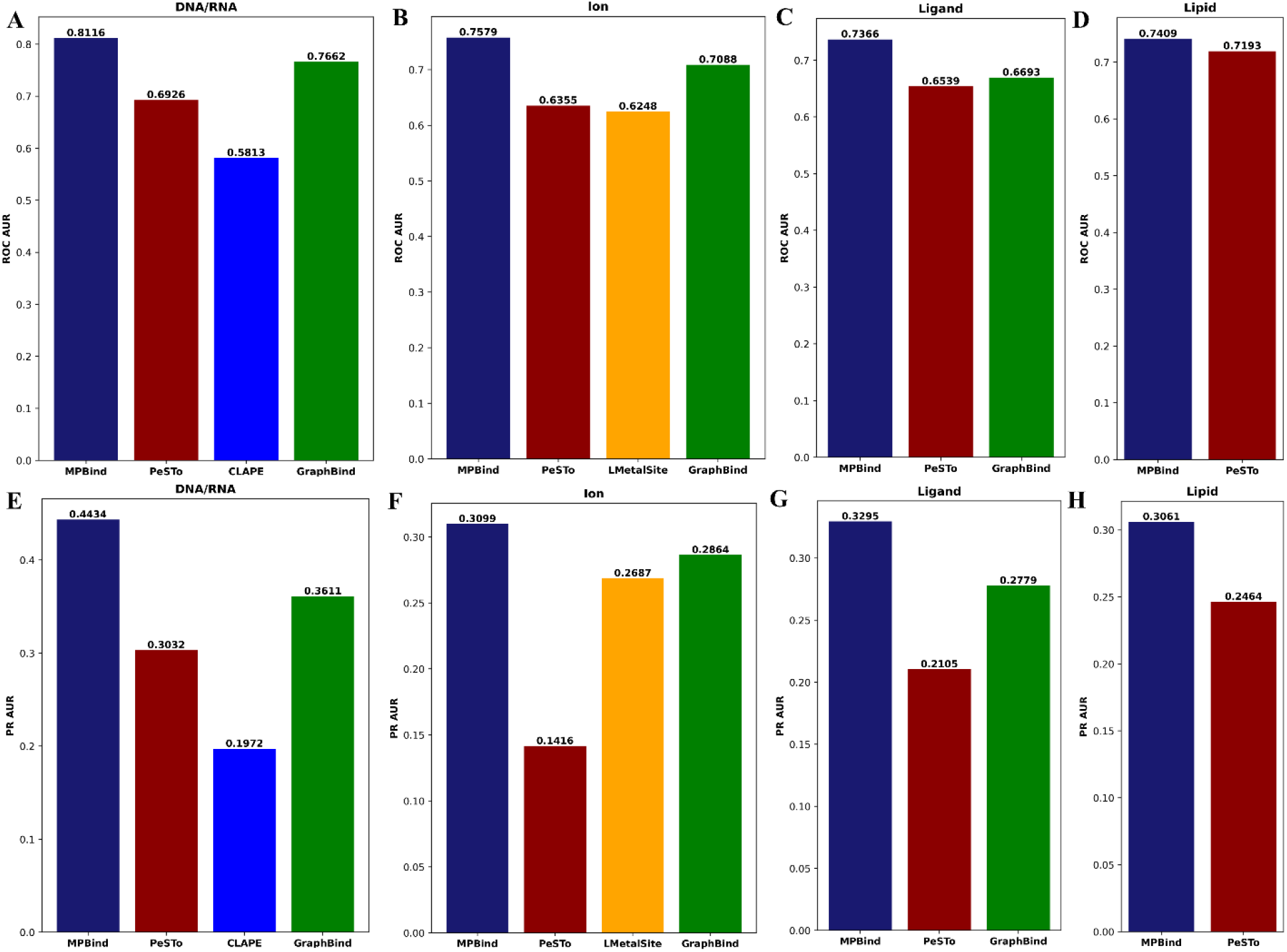
Performance comparison of MPBind with the other state-of-the-art protein-DNA/RNA, ion, ligand, and lipid binding site prediction methods on ***Test2_data***. Panels (**A-D**) present Receiver Operating Characteristic (ROC) scores, while panels (**E-F**) display Precision-Recall (PR) scores for DNA/RNA, ion, ligand, and lipid binding sites, respectively.

MPBind achieves ROC AUC scores of 0.8116, 0.7579, 0.7366, and 0.7409 for DNA/RNA, ion, ligand, and lipid binding site prediction, respectively, as shown in **Fig. 3 (A-D)**. These scores are higher than those of the other non-task-specific methods (i.e., PeSTo [11] for predicting binding site of any type) and task-specific methods (CLAPE [15] for DNA/RNA binding site prediction, LMetalSite [37] for ion binding site prediction, GraphBind [16] for DNA/RNA, ion, and ligand binding site prediction). Compared to PeSTo, the ROC AUC scores of MPBind are substantially higher for DNA/RNA, ligand, and ion binding site prediction and a little higher for lipid binding site prediction.

When considering the PR AUC metric, MPBind achieves scores of 0.4434, 0.3099, 0.3295, and 0.3061 for DNA, RNA, ion, and lipid binding site prediction, respectively, as shown in **Fig. 3 (E-H)**. These PR AUC scores are significantly higher than those of PeSTo across all binding site types, demonstrating its superior performance. Furthermore, when compared to the leading task-specific methods—CLAPE, LMetalSite, and GraphBind—MPBind also consistently outperforms them across all types of binding site predictions.

The results shows that MPBind outperforms both the general and specialized task-specific methods for binding site prediction in terms of the two critical metrics – ROC AUC and PRAUC. The results in terms of Accuracy (Acc) are detailed in supplementary **Table S2**.

Overall, the results above demonstrate that MPBind not only achieves superior results in specific types of prediction but also exhibits a remarkable level of generalization across diverse binding site types, indicating its robustness and effectiveness in accurately predicting a wide range of binding sites.

### 3.3 Five cases demonstrating MPBind’s ability to distinguish different binding interfaces

To demonstrate the general prediction capability of MPBind, we present five examples from the ***Test2_data*** dataset, each representing one of the five binding site prediction types, as shown in **Fig. 4**. The first example is the protein-protein interaction interface of the fungal halogenase RadH domain [38] that enables regioselective halogenation of various bioactive aromatic scaffolds. Figure **Fig.4(A)** depicts the protein-protein interaction within the RadH domain. When this structure is processed by MPBind, it assigns a binding site probability score between [0,1] to each residue in the right-hand chain, with the left-hand (green) chain serving as the binding partner. Among the 23 residues forming a relatively large area identified as involved in binding sites, which serve as true labels. MPBind assigns a score greater than 0.5 to 15 of these residues, identifying them as the predicted binding sites.

**Fig. 4:**
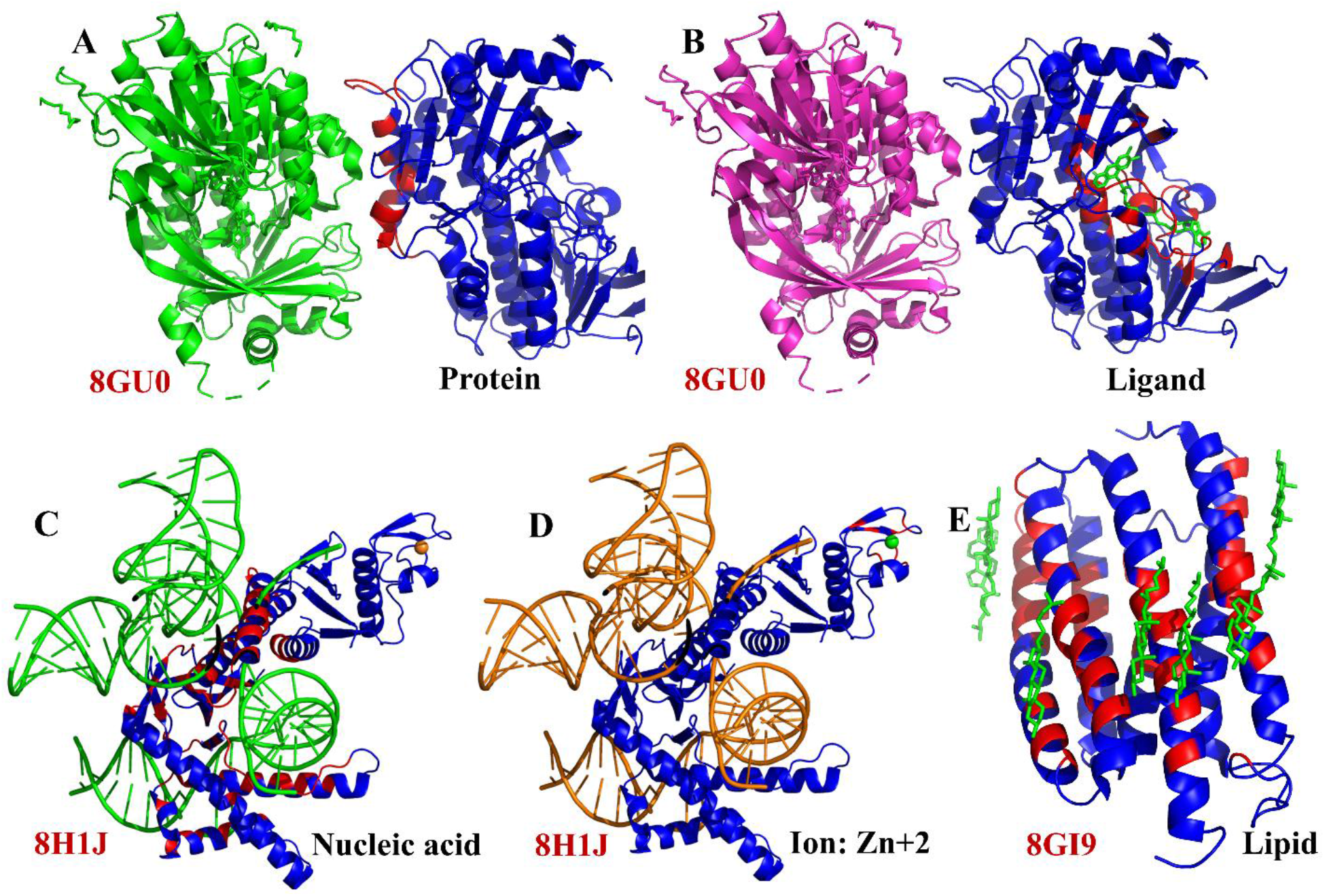
Examples of five types of binding site predictions by MPBind. The fungal halogenase RadH domain (PDB code: 8GU0) in complex with a protein **(A)** and a ligand, Flavin-Adenine Dinucleotide (FAD) **(B)**. The transposon-associated TnpB enzyme (PDB code: 8H1J) in complex with nucleic acids **(C)** and a Zn+2 ion **(D)**. Cation Channelrhodopsin (PDB code: 8GI9) in complex with multiple lipids, ChoLesteRol (CLR) **(E)**. In each subfigure, the PDB code is displayed in the bottom-left corner, the binding type is indicated in the bottom-right corner, the green region represents the partner interacting with the protein chain under consideration (for which MPBind made binding site predictions), the red region highlights the predicted binding sites, and the blue region denotes the predicted non-binding sites. In each case, some residues binding with another molecule are correctly predicted.

The second example is the same protein domain as in the first example, where it interacts with a ligand -flavin-adenine dinucleotide (FAD), colored in green, as shown in **Fig. 4(B)**. When the domain is analyzed by MPBind, it detects the ligand binding sites rather accurately. Out of the 47 residues forming a large binding pocket for FAD, MPBind assigns a prediction score above 0.5 to 31 of these residues, predicting them as the binding sites.

The third example is the *Deinococcus radiodurans* ISDra2 TnpB in complex with its cognate ωRNA and target DNA [39], shown in **Fig. 4(C)**, which involves the nucleic acid binding sites. MPBind assigns scores above 0.5 to 49 out of the 97 residues forming a large nucleotide binding area.

The fourth example involves the same protein as in the third example, as shown in **Fig. 4(D)**, which also interacts with a Zn+2 ion, represented in green. Eight residues form a binding pocket for the ion, identified as labels. MPBind predicts 7 of these 8 residues as ion binding sites with a score above 0.5.

The last example is channelrhodopsin 1 from *Hyphochytrium catenoides*, which is a light-activated channel employed for optogenetic silencing of mammalian neurons, as shown in **Fig. 4(E)**. It binds with multiple lipids. MPBind predicts 12 out of the 40 lipid-binding residues as binding sites with scores above 0.5.

The examples above show that MPBind can accurately identify some key binding residues and the location of interaction interfaces for five types of binding partners, even though it cannot predict all the binding residues at a decision threshold of 0.5. Lowering the threshold can likely predict more binding residues at the expense of a higher false positive rate.

### 3.4 Analysis of predicted binding sites in human proteome

Human proteomic data is of great interest to researchers. Here, we analyze the binding sites that predcted by MPBind for the proteins in the human proteome with annotations in UniProt [32]. We retrieved all predicted human protein structures, totaling 23,391 structures, from the AlphaFold-European Bioinformatics Institute (AF-EBI) database [31]. We selected 7,435 high-quality structures with a predicted local distance difference test (pLDDT) score of >70.0 and a predicted alignment error (PAE) of <10.0 for further analysis. The residue-annotated features for each selected structure were downloaded from the UniProt database [32].

After processing all selected structures with MPBind, we mapped the predicted binding sites to the corresponding UniProt-annotated residue-level features and calculated the proportion of UniProt entries with specific annotated features that are also predicted by MPBind to be involved in each type of binding. For instance, as shown in **Fig. 5**, more than 80% of entries of annotated non-terminal residues are predicted by MPBind to have protein-protein interaction interfaces, indicating the prevalence of non-terminal residues involved in protein-protein interactions, which is consistent with previous study [40]. In another example, more than 20% of entries with the annotated zinc finger feature are predicted to have protein-ion interaction interfaces, which is consistent with the function of zinc fingers. The results demonstrate that MPBind binding site predictions align well with the expected functional roles of the residues in the UniProt.

**Fig. 5:**
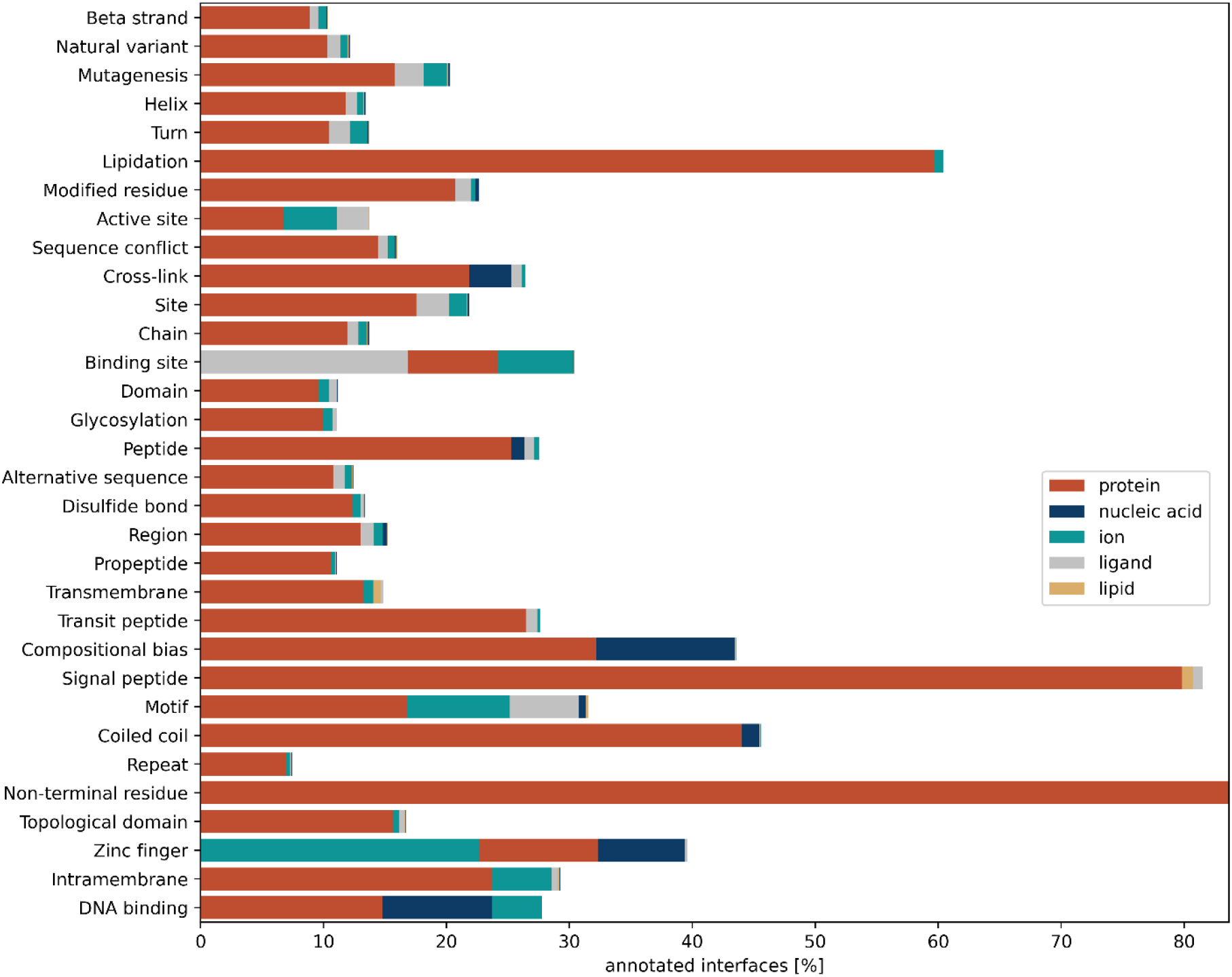
Proportion of proteins with specific annotated features in UniProt that are predicted by MPBind to have each type of binding site within their designated sequence region of the features.

We further analyze how each type of predicted binding site is related to the overall function of proteins in terms of Gene Ontology (GO) terms, including Molecular Function (MF), Biological Process (BP), and Cellular Component (CC). An interesting phenomenon, as shown in **Fig. 6** and **Fig. S1(A)**, is that the majority of entities annotated with Molecular Function (MF) GO terms are associated with protein, ligand, and ion binding sites, while fewer are linked to DNA/RNA binding sites. A similar trend is observed for Biological Process (BP) and Cellular Component (CC) Gene Ontology (GO) terms, as shown in **Fig. S1, S2**, and **S3**. Additionally, some entities annotated with MF GO terms, such as olfactory receptor activity, involve multiple binding sites, including both protein and ion binding sites simultaneously, as shown in **Fig. 6(A)** and **6(B)**. Furthermore, some entities annotated with DNA binding and RNA binding MF GO terms are primarily associated with DNA/RNA binding sites, as shown in **Fig. 6(D)**, which is also a reasonable observation. Hence, the robustness of our model is further validated through this analysis.

**Fig. 6:**
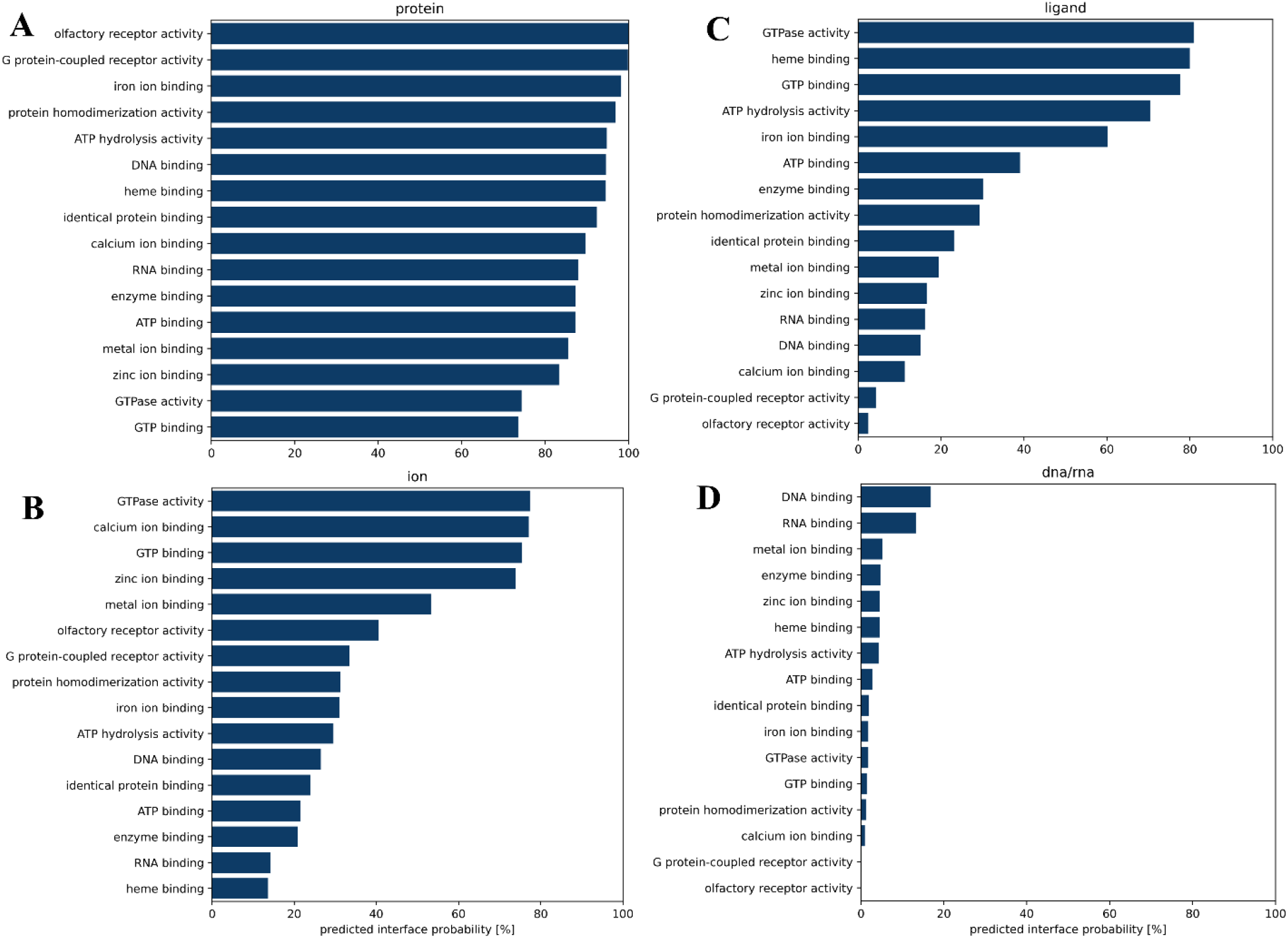
The percentage of proteins with some Gene Ontology (GO) molecular function (MF) terms in UniProt that are associated with each type of binding sites predicted by MPBind. The GO terms were ranked and selected according to the percentage. (**A**) protein binding sites, (**B**) ion binding sites, (**C**) ligand binding sites, and (**D**) DNA/RNA binding sites. For instance, over 90% of entries annotated with olfactory receptor activity are associated with the predicted protein-protein binding sites in (A), whereas almost none are linked to the predicted DNA/RNA-binding site interfaces in (D).

### 3.5 An ablation study of different features in MPBind

We investigated the contribution of each type of node feature to the performance of MPBind on the ***Test2_data*** dataset. Starting from the four types of features (ProtTrans sequence embedding, DSSP features, Atomic Residue features, and Geometric Residue features) with ROC AUC = 0.77 and PR AUC = 0.54 (the first row in **Table 1**), we added one of the two kinds of embeddings generated by ProstT5 (ProstT5(AA): amino acid sequence embedding and ProstT5(3Di: ProstT5 structural alphabet embedding), respectively. Adding ProstT5(AA) substantially increased the ROC AUC and PR AUC to 0.84 and 0.62 (the second row in **Table 1**), while adding ProstT(3Di) did not affect ROC AUC and only slightly increased PR AUC to 0.55 (the third row in **Table 1**). Therefore, we chose to only use ProstT5(AA) features with the other four types of features as the start point in the following three experiments. We then tried to delete one of the three types (DSSP features, Atomic Residue features, and Geometric Residue features) in the three separate experiments (the 4^th^, 5^th^, and 6^th^ rows in **Table 1**). Deleting any one of them decreased the performance, indicating that each of them is useful, while the impact of the DSSP features is higher than the Atomic Residue features, and the Geometric Residue features are least important. The ablation study demonstrates that using ProtTrans, ProstT5 (AA), DSSP, Atomic Residue, and Geometric Residue features together yields the best results. The combination of the five types of features is used in the final version of MPBind.

**Table 1.** Contribution of different node feature types to the performance of MPBind on the ***Test2_data*** dataset in terms of ROC AUC and PR AUC.

| ProtTrans | ProstT5 (AA) | ProstT5 (3Di) | DSSP | Atomic Residue | Geometric Residue | ROC AUC | PR AUC |
| --- | --- | --- | --- | --- | --- | --- | --- |
| ✓ | ✗ | ✗ | ✓ | ✓ | ✓ | 0.77 | 0.54 |
| ✓ | ✓ | ✗ | ✓ | ✓ | ✓ | **0.84** | **0.62** |
| ✓ | ✗ | ✓ | ✓ | ✓ | ✓ | 0.77 | 0.55 |
| ✓ | ✓ | ✗ | ✗ | ✓ | ✓ | 0.78 | 0.57 |
| ✓ | ✓ | ✗ | ✓ | ✗ | ✓ | 0.80 | 0.59 |
| ✓ | ✓ | ✗ | ✓ | ✓ | ✗ | 0.81 | 0.60 |

✓ = feature included ✗ = feature removed

## Conclusion

In this study, we developed a new multitasking method (MPBind) of using protein language models and an equivariant graph neural network to integrate both protein sequence and structure data to predict five types of protein binding sites. A graph with some new node and edge features are designed to capture the sequence, structural, and geometrical features of input proteins to be processed by MPBind. MPBind can not only predict a broad spectrum of protein binding sites interacting with proteins, DNA, RNA, ligands, ions, and lipids, but also outperforms other general or task-specific binding site prediction methods. Additional analysis of MPBind predicted binding sites for the human proteins show that they are consistent with the annotated functional features of the proteins in the UniProt. The results demonstrate that MPBind can be used to predict the binding sites of unannotated proteins and facilitate their functional and structural study.

## Supporting information

Supplemental for MPBind

## Data availability

The original protein structures used in this work can be downloaded from the PDB by the script provided at our GitHub repository: https://github.com/jianlin-cheng/MPBind. The scripts for generating training and test datasets from the structures are also provided at the GitHub repository.

## Code availability

The source code of MPBind is available at the GitHub repository: https://github.com/jianlin-cheng/MPBind.

## Acknowledgments

This work was supported in part by the grants from the USA National Science Foundation [NSF grant numbers: DBI2308699, CCF2343612].

