## Supplemental for MPBind for "MPBind: Multitask Protein Binding Site Prediction by Protein Language Models and Equivariant Graph Neural Networks"

**Table.S1:** Performance comparison of MPBind and two protein-protein binding site prediction methods across two test datasets. Bold fond denotes the best result.

| **Method** | ***Test1_data*** | ***Test2_data*** |
|---|---|---|
| | **Acc** | **Acc** |
| MPBind | **0.8468** | **0.8271** |
| PeSTo | 0.7905 | 0.7559 |
| ScanNet | --- | 0.7902 |

**Table.S2:** Performance comparison of MPBind and state-of-the-art protein-DNA/RNA, ion, ligand, and lipid binding site prediction methods on ***Test2_data*** dataset.

| **Binding site type** | **Method** | ***Test2_data*** |
|---|---|---|
| | | **Acc** |
| **DNA/RNA** | MPBind | **0.9902** |
| | PeSTo | 0.9372 |
| | CLAPE | 0.8825 |
| | GraphBind | 0.8773 |
| **Ion** | MPBind | **0.9875** |
| | PeSTo | 0.9874 |
| | LMetalSite ($Ca^{2+}$, $Mg^{2+}$, $Mn^{2+}$, $Zn^{2+}$) | 0.9673 |
| | GraphBind ($Ca^{2+}$, $Mg^{2+}$, $Mn^{2+}$) | 0.9666 |
| **Ligand** | MPBind | **0.9761** |
| | PeSTo | 0.9744 |
| | GraphBind (ATP, HEME) | 0.9203 |
| **Lipid** | MPBind | **0.9972** |
| | PeSTo | 0.9505 |

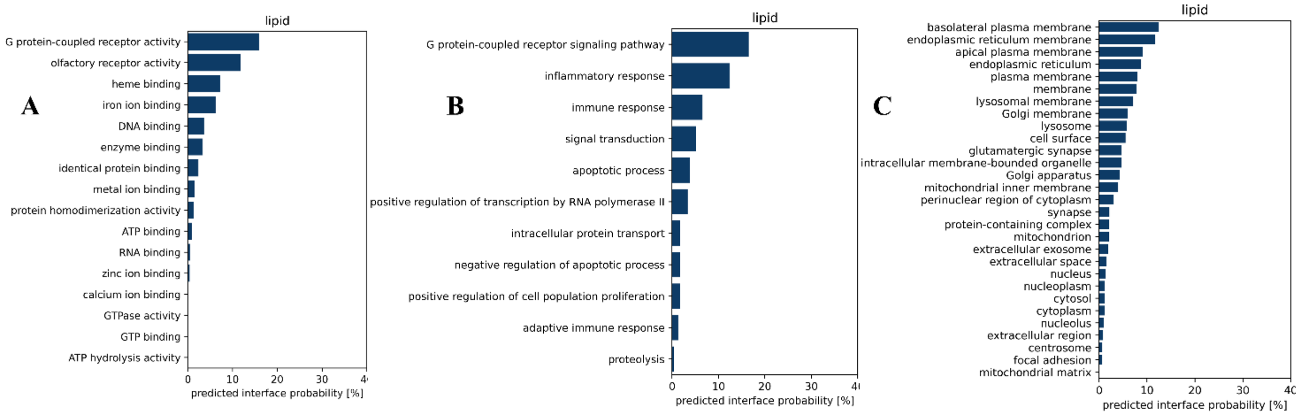


**Fig. S1:** The percentage of proteins with some Gene Ontology (GO) terms from UniProt that are associated with lipid binding sites predicted by MPBind. The GO terms were ranked and selected according to the percentage. (**A**) molecular function (MF) terms, (**B**) biological process (BP) terms, and (**C**) cellular component (CC) terms.

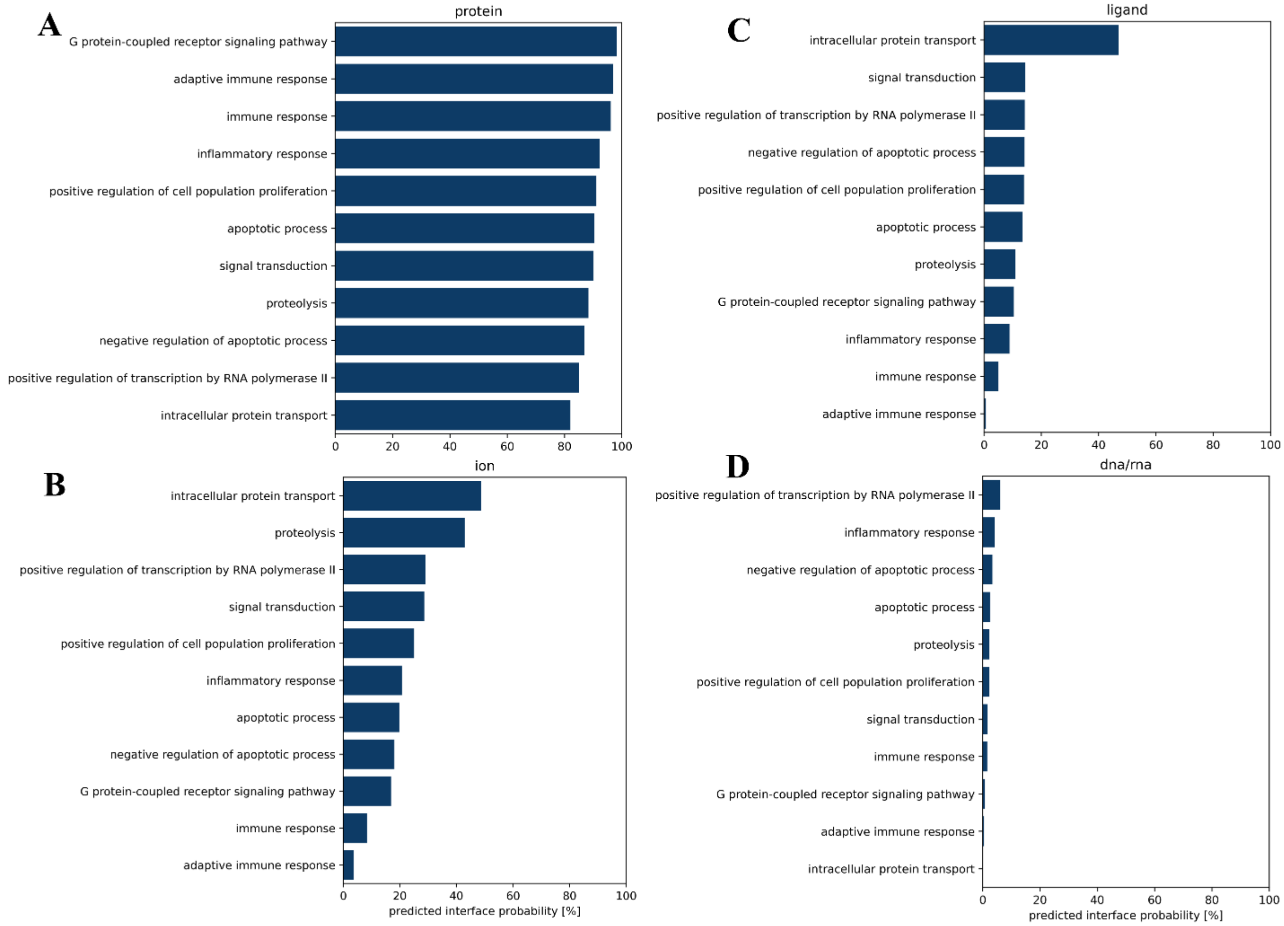


**Fig. S2:** The percentage of proteins with some biological process GO terms from UniProt that are associated with four different types of binding sites predicted by MPBind. The GO terms were ranked and selected according to the percentage. (**A**) protein binding sites, (**B**) ion binding sites, (**C**) ligand binding sites, and (**D**) DNA/RNA binding sites.

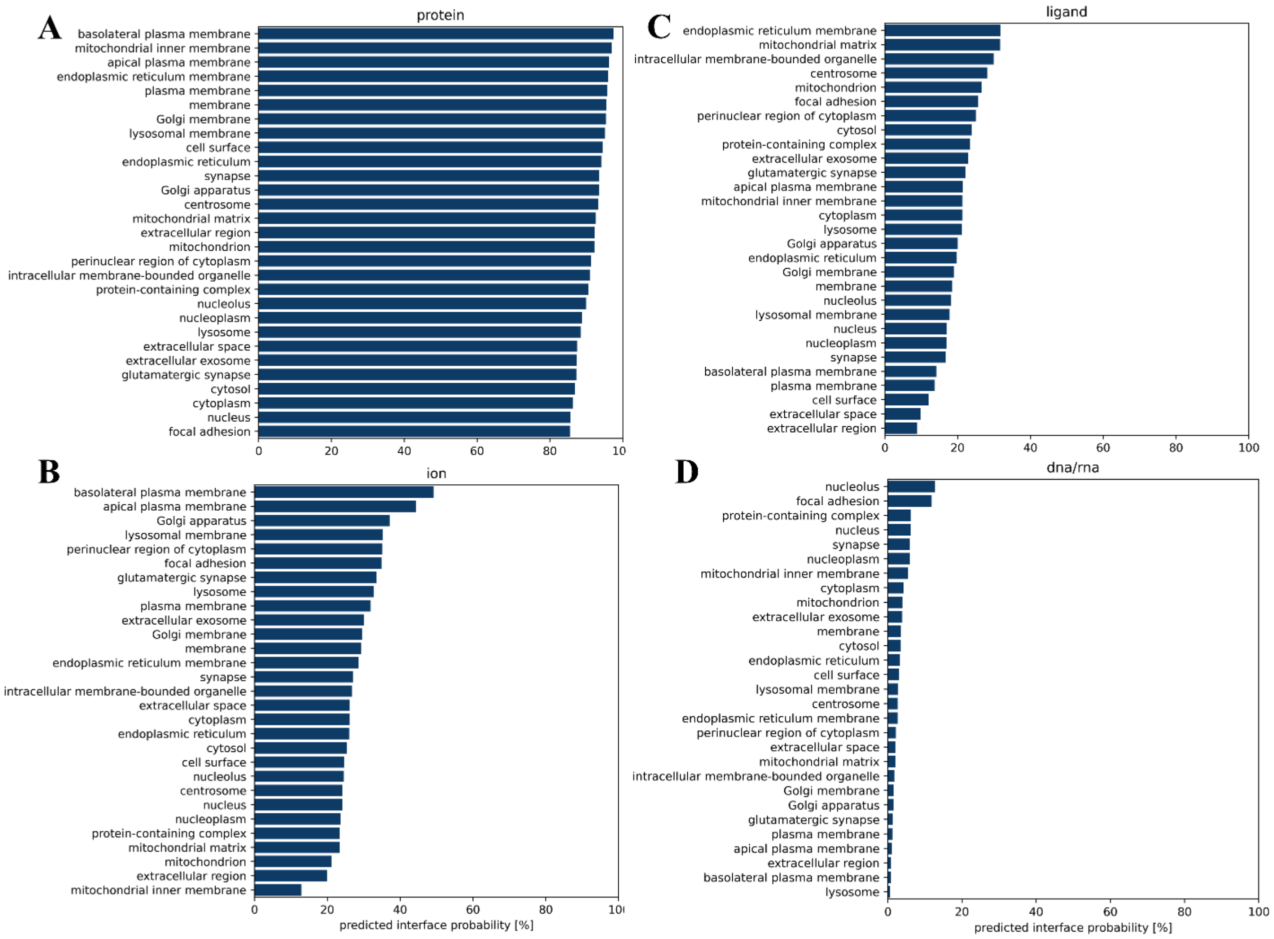


**Fig. S3:** The percentage of proteins with some cellular component GO terms from UniProt that are associated with distinct types of binding sites predicted by MPBind. The GO terms were ranked and selected according to the percentage. (**A**) protein binding sites, (**B**) ion binding sites, (**C**) ligand binding sites, and (**D**) DNA/RNA binding sites.
